## Supplemental Figures for "Tuberculosis susceptibility in genetically diverse mice reveals functional diversity of neutrophils"

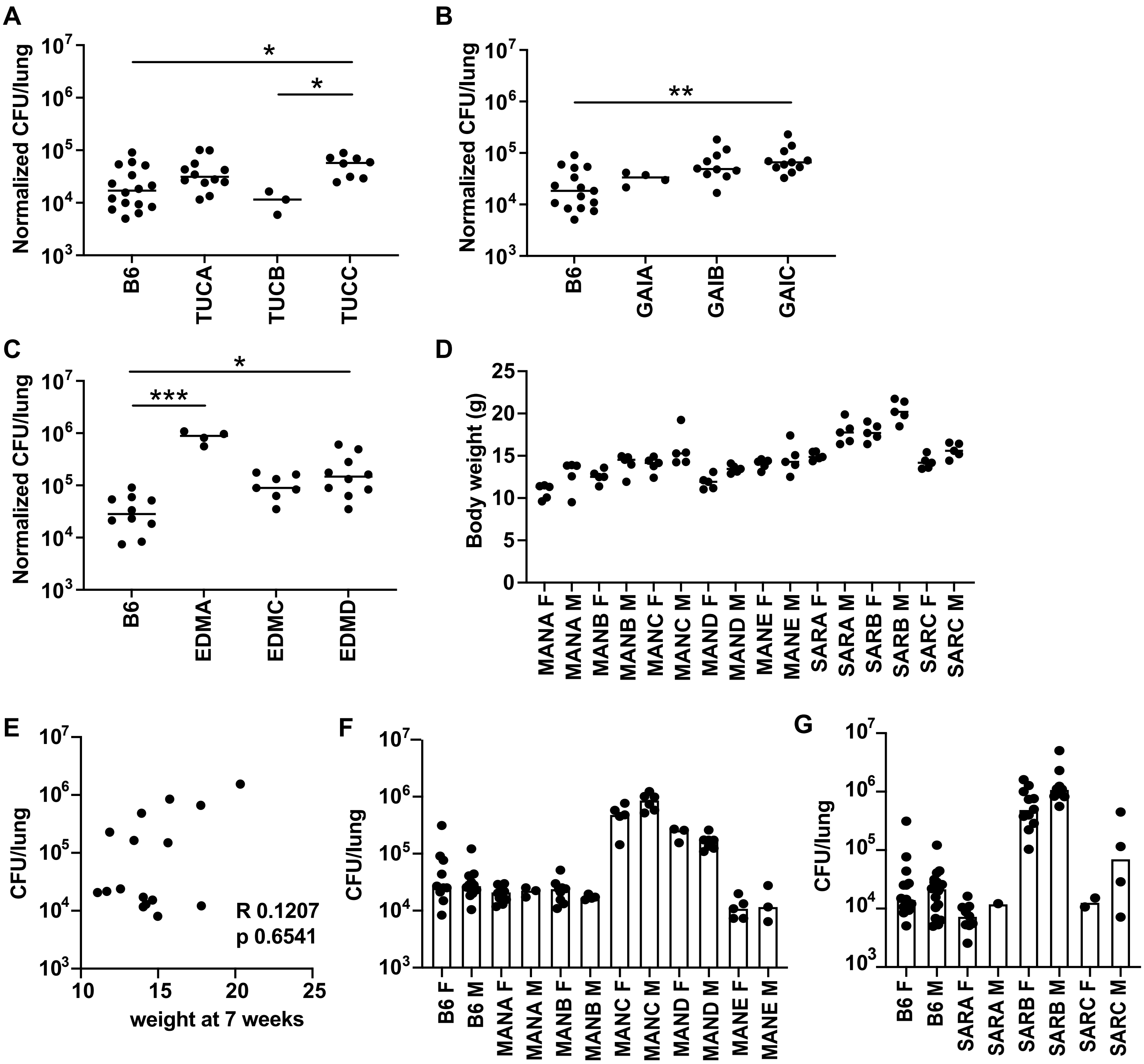


**Figure 1 – figure supplement 1. Variation in susceptibility to infection in wild mice does not depend on mouse weight or sex.** CFU in the lungs of mice from (A) Tucson (B) Gainesville and (C) Edmonton at 21 days post infection were enumerated by CFU. (D) Body weight of adult male (M) and female (F) mice at age 7 weeks of age. (E) Spearman correlation analysis of CFU in the lungs vs weight at 7 weeks. (F,G) CFU data from the lungs of mice at 21d post infection broken down by both sex and genotype. CFU data was normalized to CFU enumeration on day 1 to correct for variation in inoculum dose between experimental days. Data represent of 2-3 independent experiments, except for a single experiment for the TUCC, EDMA, EDMC and EDMD lines. Each dot represents a mouse. *p<0.05, **p<0.01, ***p<0.001 by Mann Whitney U test.

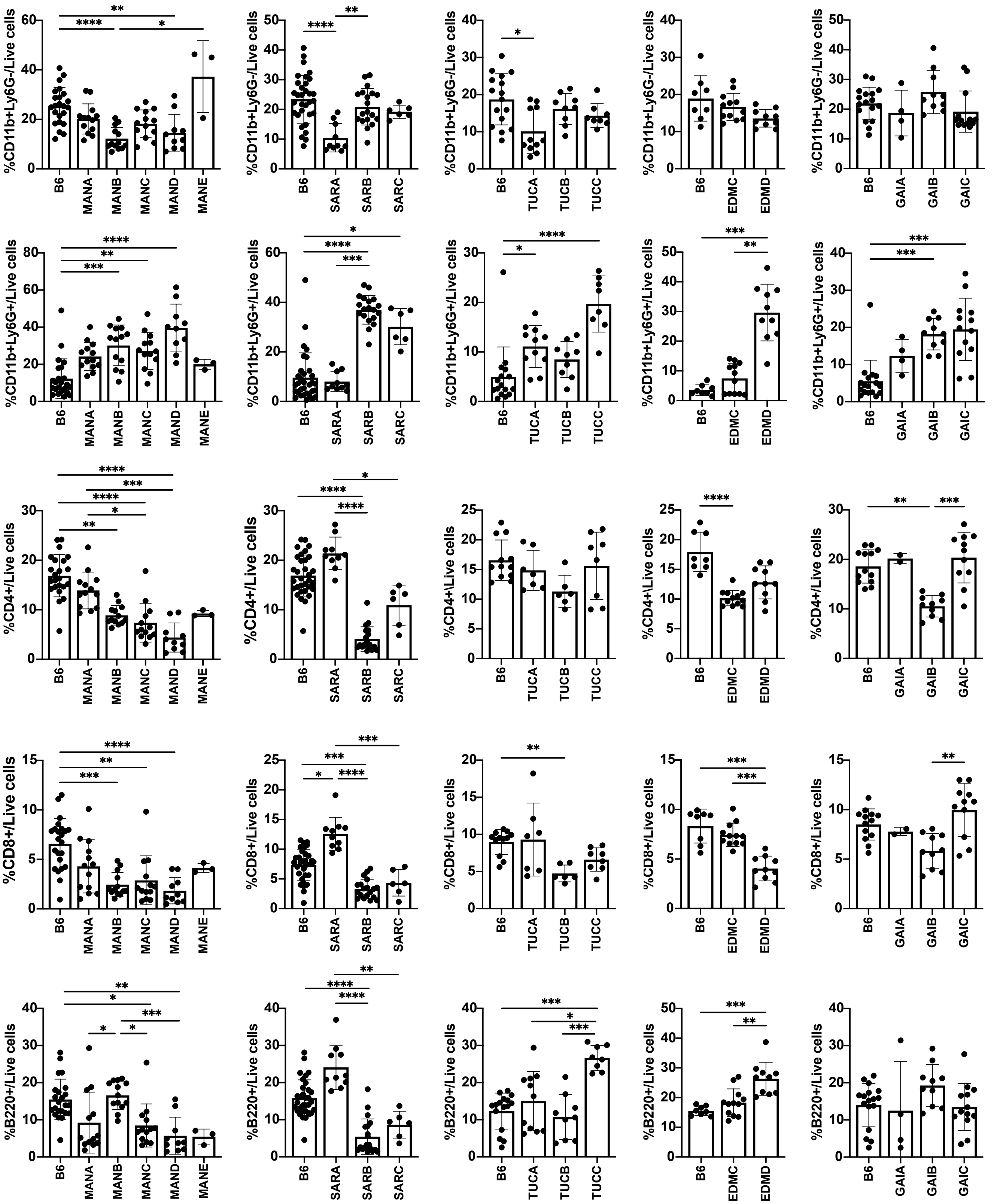


**Figure 2 – figure supplement 1. Cell percentages from flow cytometry-based analysis in the lungs of infected wild mice relative to B6.** Percentages of live cell populations were enumerated at 21 days post infection with *Mtb.* Each circle represents an individual mouse. Data is representative of 2-6 independent experiments for each line. The $p$ values were determined using a Kruskal-Wallis ANOVA. $^{*}p < 0.05$, $^{**}p < 0.005$, $^{***}p < 0.0001$, $^{****}p < 0.0001$.

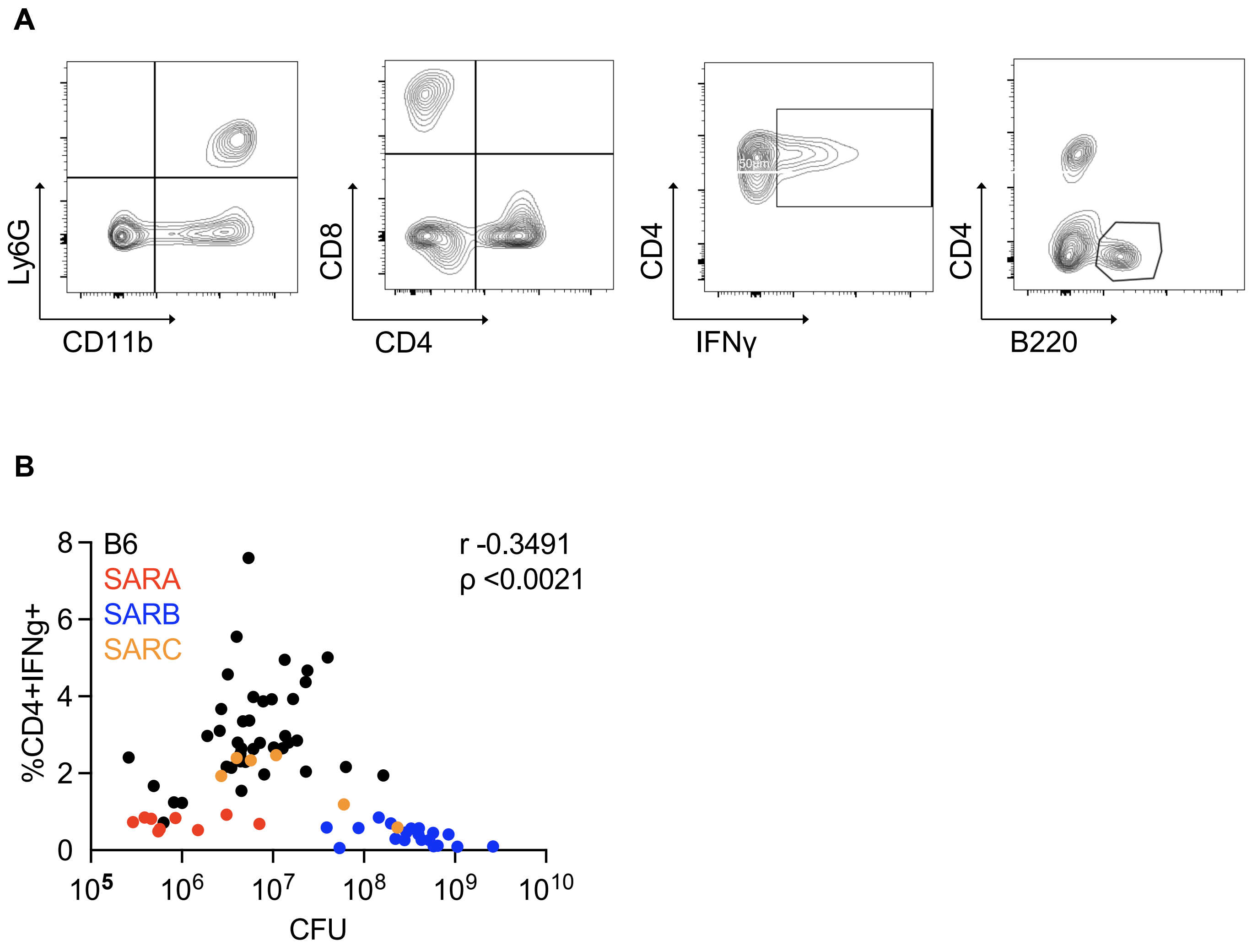


**Figure 2 – figure supplement 2. The proportion of IFN-γ producing CD4 T cells in the lung does not correlate with bacterial burden.** (A) Gating strategy for individual cell types in the lung. All cell populations were gated on live cells. CD4 and CD8 cell populations were identified from CD3+ cells, followed by IFN-γ staining. Neutrophils were identified as CD11b+Ly6G+ and macrophages/monocytes as CD11b+Ly6G-. (B) Spearman correlation of CFU vs CD4+IFN-γ+ cells in the lungs of B6 or SARA, SARB and SARC mouse lines infected with *Mtb* enumerated at 21 days post infection. Data represent at least 2 independent experiments per line. The *p* value was determined using two-tailed Spearman correlations.

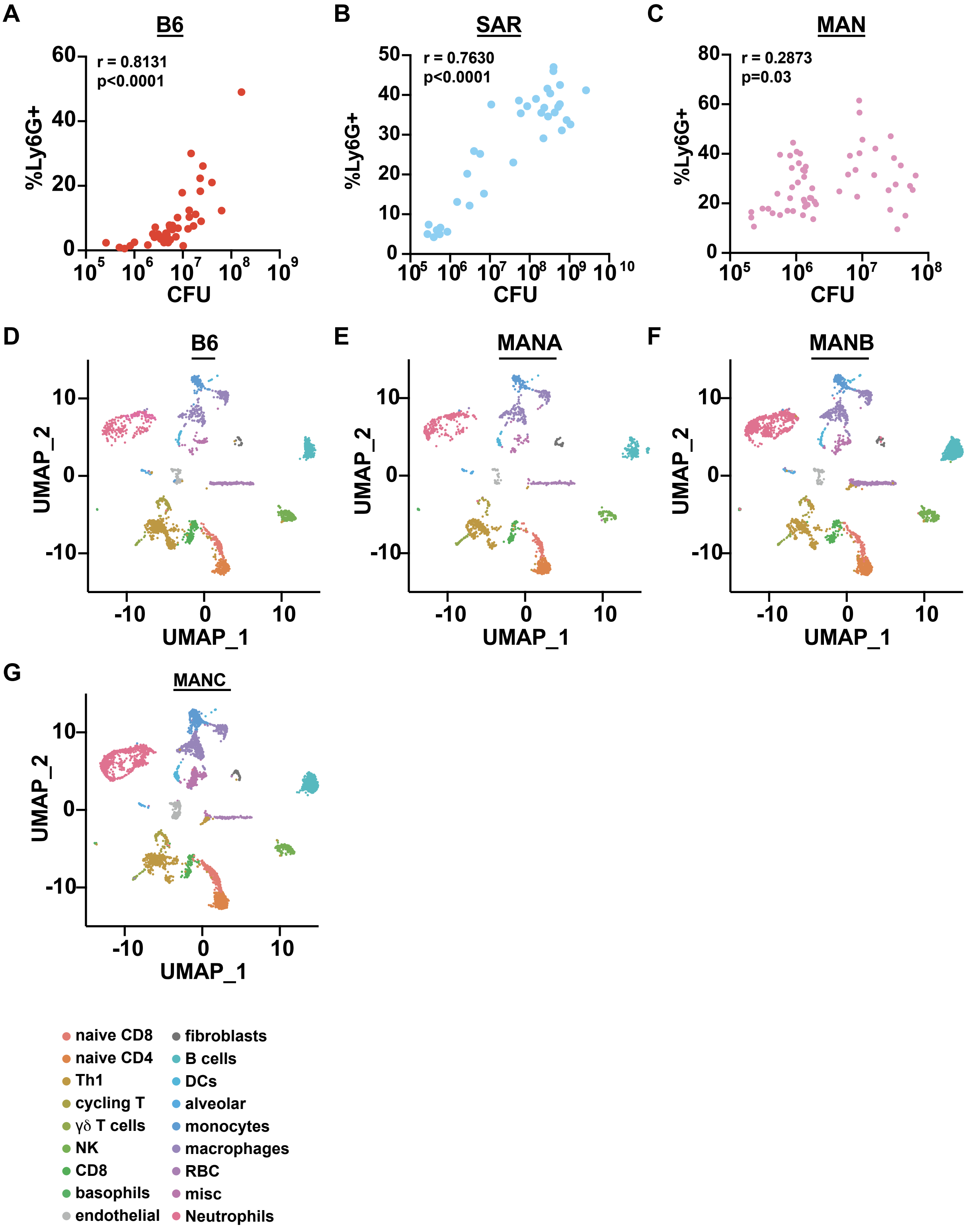


**Figure 2 – figure supplement 3. Immune cell composition in the lungs of infected mice.** Spearman correlations between bacterial burden and presence of neutrophils (CD11b+Ly6G+) in the lung of infected B6 (A), Saratoga (B) and Manaus (C) mice. (D-G) UMAP plots of single cells isolated from the lungs of *Mtb* infected mice at 21 days post infection. (D) B6, (E) MANA, (F) MANB, (G) MANC. (H-J) Gene expression analysis of lung neutrophils scored according to gene signatures identified by Xie *et al*[61].

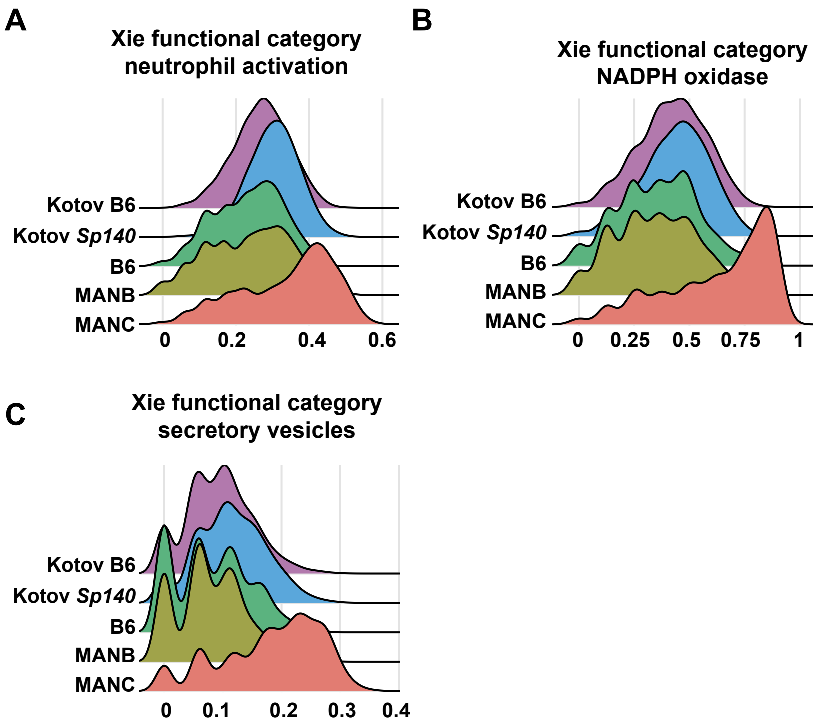


**Figure 4 – figure supplement 1. Immune cell composition in the lungs of infected mice.** (A-C) Gene expression analysis of lung neutrophils scored according to gene signatures identified by Xie *et al*[61].

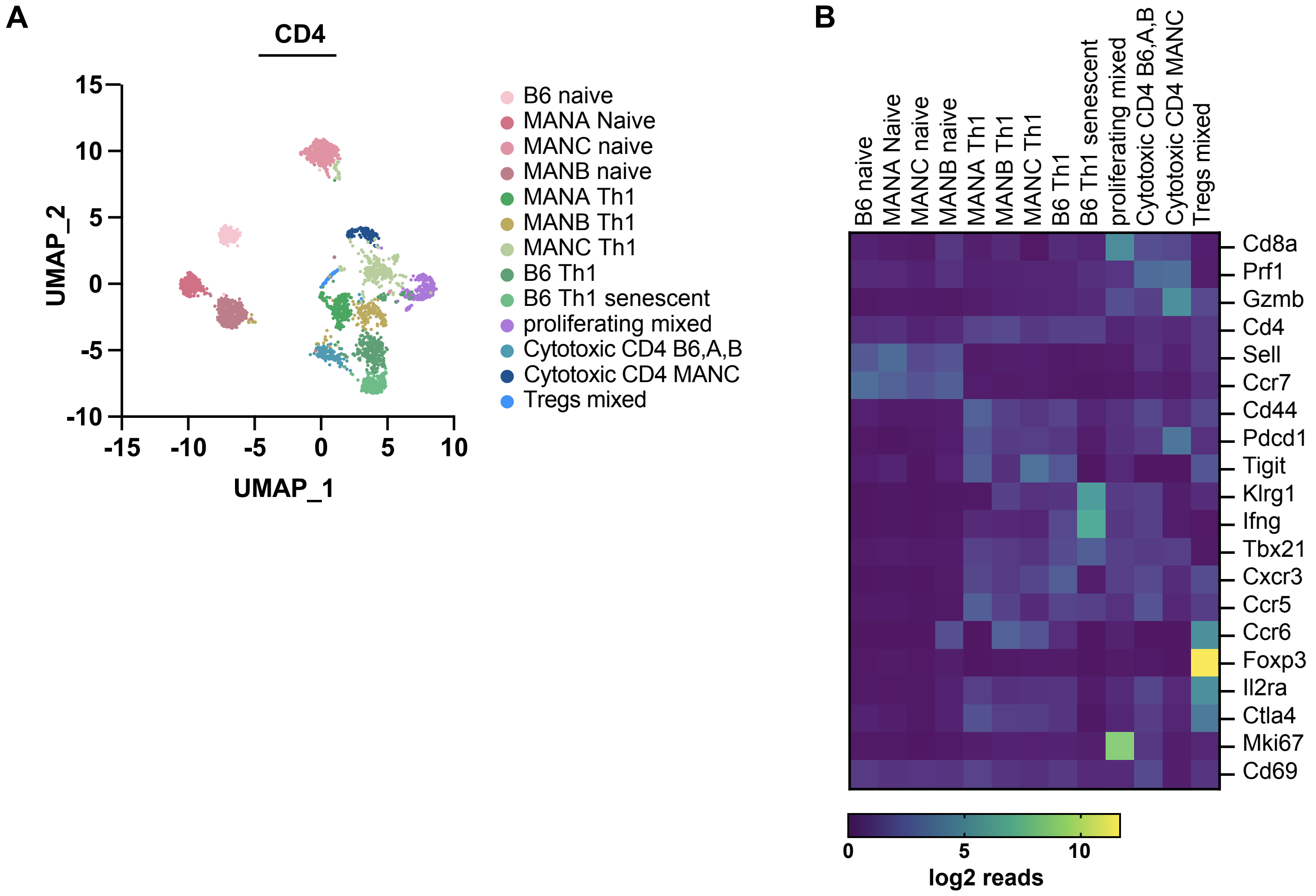


**Figure 4 – figure supplement 2. T cell subsets and gene expression in wild-derived mice.** (A) UMAP analysis of CD4 T cell subsets in the lung from scRNAseq expression data of Manaus and B6 mice. (B) Heatmap of scRNAseq expression data of T cell clusters from individual Manaus lines and clusters from mixed mouse lines. Significance calculation is students t-test, $p < 0.05$.

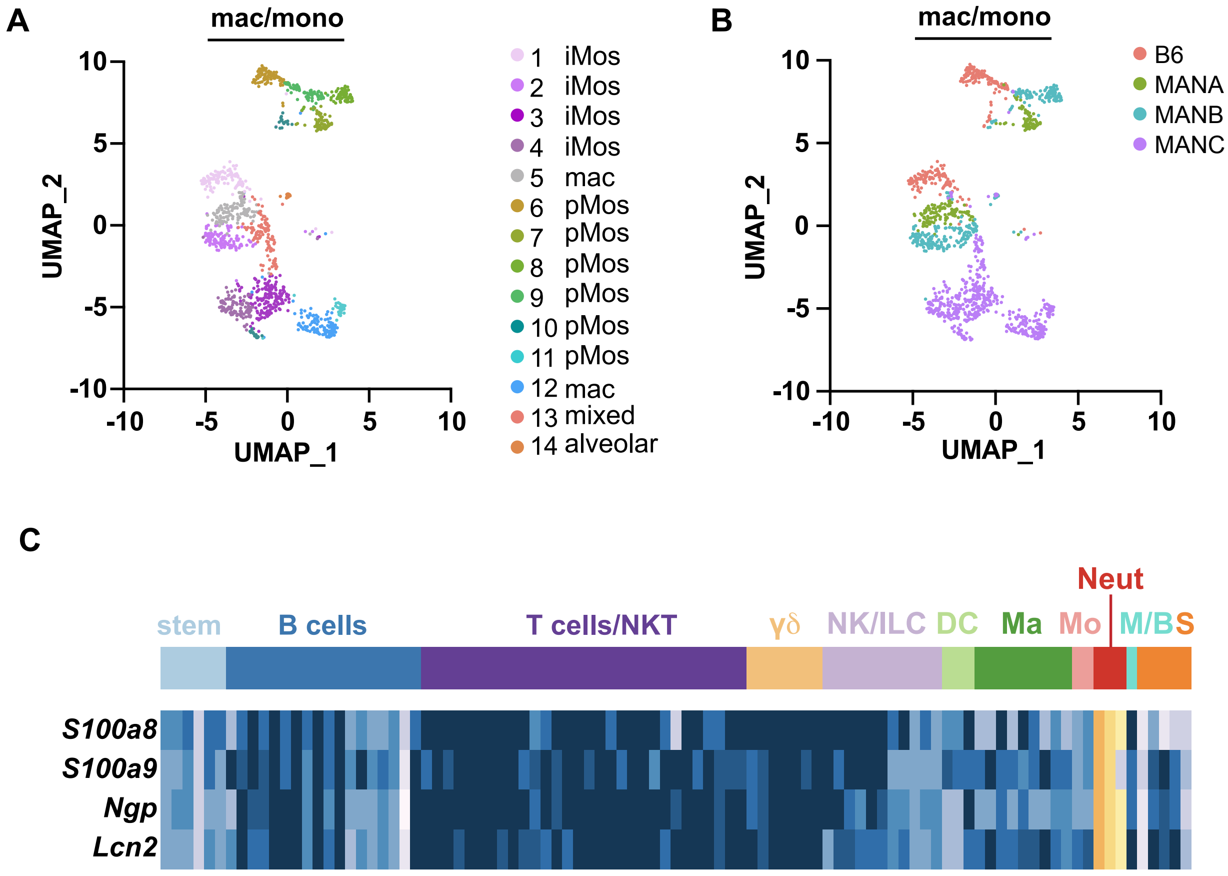


**Figure 6 – figure supplement 1. Gene expression across immune cells.** (A) Unsupervised clustering of CD11b+Ly6G- monocyte/macrophage-like cells in the lung based on scRNAseq gene expression color coded by cluster (A) or genotype (B). (C) Expression of indicated genes across immune cell types analyzed using Immunological Genome Project (ImmGen) MyGeneset application (http://rstats.immgen.org/MyGeneSet_New/index.html). Inflammatory monocytes (iMos); patrolling monocytes (pMos), stem cells (stem); B cells; T cells/NK T cells (NKT); γδ T cells (γδ); Natural Killer/innate lymphoid cells (NK/ILC); dendritic cells (DC); macrophages (Ma); monocytes (Mo); neutrophils (Neut); Mast Cells/Basophils (M/B); Stromal Cells (S).
